# A-Intercalated Cell Dysfunction Disrupts Renal Epithelial–Immune Balance and Impairs Host Defense During UTI

**DOI:** 10.64898/2026.02.10.705112

**Authors:** Forough Chelangarimiyandoab, Kristina McNaughton, Grace Essuman, Emmanuelle Cordat

## Abstract

Intercalated cells (ICs) of the renal collecting duct are traditionally recognized for their role in acid–base homeostasis, but growing evidence suggests they also participate in innate immune defense. Although ICs have been implicated in renal antimicrobial function, their specific role in coordinating immune responses during urinary tract infection (UTI) remains unclear. Using Ae1 R607H knock-in mice, a distal renal tubular acidosis (dRTA) model with A-intercalated cell (A-IC) dysfunction, we examined the renal response to uropathogenic *Escherichia coli* (UPEC). Mice with A-IC dysfunction exhibited higher bacterial loads 24 h post-infection and increased renal expression of antimicrobial peptides lipocalin-2 (*Lcn2*), galectin-3 (*Lgals3*), and cathelicidin-related antimicrobial peptide (*Camp*). Pro-inflammatory cytokines interleukin-6 (IL-6) and interleukin-1β (IL-1β) were elevated at both transcript and protein levels, whereas tumor necrosis factor-α (TNF-α) increased only at the protein level. Interleukin-10 (IL-10) showed a modest rise in mRNA. Chemokines C-X-C motif chemokine ligand 2 (*Cxcl2*) and C-C motif chemokine ligand 2 (*Ccl2*) were also upregulated, accompanied by excessive neutrophil infiltration and a marked shift in renal myeloid-cell composition. A-IC dysfunction therefore disrupts epithelial–immune homeostasis, resulting in exaggerated inflammation and impaired immune resolution. These findings identify A-ICs as essential epithelial immunomodulators that integrate antimicrobial defense, cytokine regulation, and immune-cell recruitment during UTI.

## INTRODUCTION

Urinary tract infections (UTIs) are among the most common bacterial infections worldwide, disproportionately affecting women ^1^. About 60% of women will experience at least one UTI in their lifetime, and nearly 25% will suffer from recurrent infections ^2,3^. The rapid increase in antibiotic resistance underscores the urgency of developing alternative treatments for UTIs ^4^. These infections can impact different segments of the urinary tract, categorized as urethritis, cystitis, and pyelonephritis ^5^. UTIs can be caused by various pathogens, including Gram-negative and Gram-positive bacteria, as well as fungi, with uropathogenic *Escherichia coli* (UPEC) being the most common cause ^6^.

As the UPEC ascends through the urinary tract, the renal collecting duct (CD) becomes the first segment of the kidney to encounter the pathogen ^7^. The CD is composed of two primary cell types: principal cells (PCs) and intercalated cells (ICs) ^8^. PCs are involved in sodium and water reabsorption, whereas ICs regulate acid-base homeostasis and participate in potassium and ammonia transport ^8,9^. ICs are further classified into three subtypes: A-IC, B-IC, and non-A, non-B IC ^10^. Beyond their role in electrolyte and pH regulation, ICs have emerged as key players in the kidney’s innate immune defense ^7,11^. Uropathogenic bacteria preferentially adhere to their apical surface^12^. ICs can directly engage invading pathogens through phagocytosis and acidification of bacteria-containing compartments ^12^. Additionally, ICs express antimicrobial peptides (AMPs) and pattern recognition receptors such as Toll-like receptor 4 (TLR4) and purinergic receptor P2Y, G-protein coupled, 14 (P2Y14), enabling detection of pathogens and tissue damage ^12–16^. Studies show that IC deficiency increases susceptibility to bacteriuria and pyelonephritis, highlighting their protective role ^12,17^. ICs also mediate inflammation by recruiting immune cells via chemokine release, and transcriptomic data reveal a coordinated innate immune gene program in ICs ^15,18,19.^ Distal renal tubular acidosis (dRTA) is a genetic disorder caused by defective ICs, resulting in systemic metabolic acidosis and impaired urinary acidification ^9,10,19–21^. While dRTA can arise from variations in several genes, this study focuses on solute carrier family 4 member 1 (*SLC4A1*), which encodes the anion exchanger 1 (AE1) protein expressed in A-ICs ^22–24^. Kidney biopsies from dRTA patients, as well as studies using R607H knock-in (KI) mouse model—which mirrors the dominant AE1 R589H variation found in human dRTA—demonstrate a cortical reduction in A-IC abundance, along with a broader impairment of A-IC phenotype ^25,26^. Although medullary A-ICs remain present, they exhibit decreased expression of AE1 and B1 subunit of vacuolar H^+^-ATPase (V-ATPase B1), indicating functional compromise of this cell type ^27^. As Ae1 R607H KI mice exhibit an alkaline urine and metabolic acidosis after an acid challenge due to A-IC dysfunction, they provide an appropriate model to examine the role of A-IC in bacterial clearance.

Although limited clinical observations suggest that dRTA patients may be more susceptible to UTIs, the immune mechanisms underlying this susceptibility remain largely unexplored ^28^. Given the established A-IC dysfunction in the Ae1 R607H KI mouse, we hypothesized that impaired A-IC function compromises renal innate immunity, leading to delayed bacterial clearance and prolonged inflammation during UTI. This study aimed to characterize the immune response to UPEC in the Ae1 R607H KI mice, shedding light on the role of ICs in infection resolution and local inflammatory regulation.

## MATERIALS AND METHODS

### Animal Model

All animal procedures were conducted in accordance with the guidelines of the Canadian Council on Animal Care (CCAC) and were approved by the Animal Care and Use Committee (ACUC) at the University of Alberta under Animal Use Protocol (AUP No. 1277). Female wild-type (WT) and Ae1 R607H homozygous (HO) dRTA KI mice aged 8–10 weeks were used for all experiments. The Ae1 R607H mice have previously been characterized ^27^. All animals were housed in Health Sciences Laboratory Animal Services (HSLAS) Biosafety Level 2 under a 12-hour light/dark cycle with *ad libitum* access to water and standard chow (PicoLab Mouse Diet 20, no. 5058).

### UTI Induction

Mice were deprived of drinking water for 30 minutes prior to anesthesia to minimize immediate voiding post-inoculation. Before anesthesia, mice were placed in a sterile container, and a gentle massage of the caudal abdomen was performed to induce urination and ensure an empty bladder ^29^. Mice were then anesthetized with isoflurane, and upon reaching a surgical plane of anesthesia, the vulva and perineum were swabbed with 70% ethanol to reduce the risk of external contamination. A sterile 1 cc syringe attached to a lubricated, sterile catheter/cannula was inserted into the urethra, and 50 µL of either bacterial suspension (20–100 × 10^6^ colony-forming units (CFUs)) or sterile phosphate-buffered saline (PBS) was slowly instilled into the bladder ^30^. Mice were maintained under anesthesia for up to 20 minutes post-inoculation to prevent immediate voiding, then allowed to recover on a warming pad under close observation before being returned to their home cages.

### Statistical Analysis

Statistical analyses were performed using GraphPad Prism (v10.4.1). When appropriate, data were normalized to the average value of the WT PBS group before statistical testing. Data were assessed for normality, and outliers identified by the software were excluded. Depending on the distribution and experimental design, comparisons between two groups were made using either an unpaired two-tailed Student’s *t*-test or the Mann–Whitney *U* test. For multiple group comparisons, two-way ANOVA followed by Tukey’s multiple-comparisons test was used. For flow cytometry experiments where specific groups were directly compared, unpaired two-tailed *t*-tests were applied. A *p*-value of less than 0.05 was considered statistically significant. Results are expressed as mean ± standard error of the mean (S.E.M.).

**Additional detailed experimental procedures are provided in the *Supplementary Material***.

## RESULTS

### A-IC dysfunction impairs bacterial clearance in dRTA mice

ICs play an important role in renal defense against pathogens through acid-base regulation and secretion of antimicrobial factors ^7,8,12,14^. To determine whether **A-IC dysfunction** compromises host resistance to infection, we first examined the expression of A-IC–associated markers in WT or HO mice transurethrally catheterized with sterile PBS or UPEC. The UPEC strain used in this study was originally isolated from a female patient with pyelonephritis and expresses the type 1 fimbrial adhesin FimH but lacks the hemolysin HlyA virulence factor. This FimH^+^/HlyA^−^, non-hemolytic phenotype is typical of UPEC strains that adhere to urinary tract epithelia and elicit robust immune activation without extensive cytotoxicity ^31–33^.

Consistent with previous characterization ^27^, quantitative reverse transcription polymerase chain reaction (qRT-PCR) analysis confirmed reduced expression of IC markers—AE1 (*Slc4a1*), V-ATPase B1 (*Atp6v1b1*), and carbonic anhydrase 2 (*Car2*)—in HO mice compared with WT controls under both PBS and UPEC conditions (**Figure 1**). UPEC infection did not alter gene expression of these 3 markers compared to PBS.

**Figure 1.**
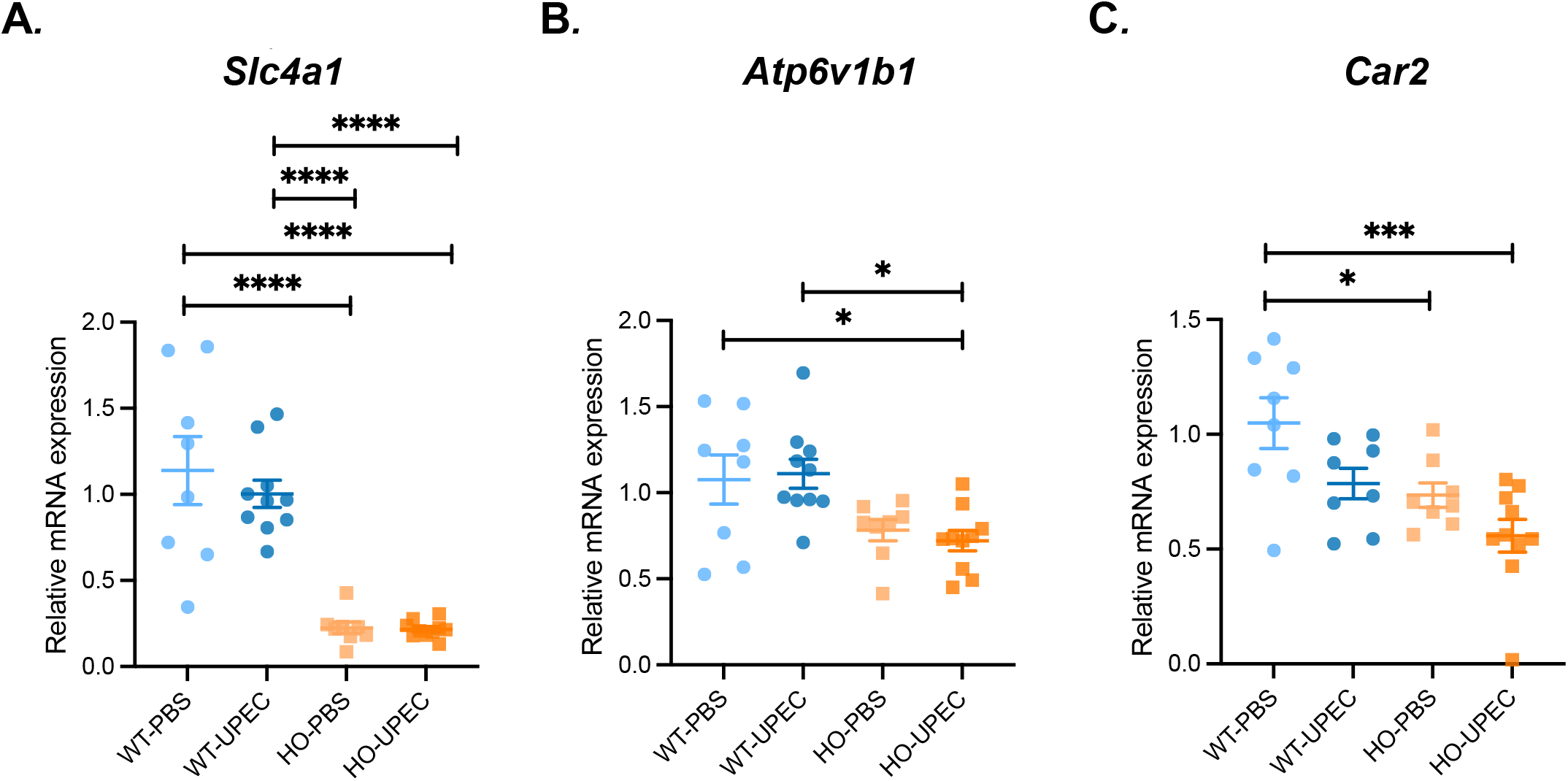
Reduced expression of A-IC markers in Ae1 R607H KI mice. Relative mRNA expression of A-IC markers *Slc4a1* (A), *Atp6v1b1* (B), and *Car2* (C) in kidneys from WT or HO Ae1 R607H KI mice injected with PBS or UPEC, measured by qRT-PCR. Data represent relative mRNA expression normalized to *Rplp0* and expressed relative to WT kidneys exposed to PBS using the 2^-ΔΔCT method. Data correspond to mean ± SEM, and each symbol indicates one animal. Statistical analysis was performed using two-way ANOVA followed by Tukey’s multiple-comparisons test (\**p* < 0.05; \*\*\**p* < 0.001; \*\*\*\**p* < 0.0001).

To evaluate the impact of A-IC dysfunction on infection control, CFUs were quantified in bladder and kidney tissue homogenates after UPEC inoculation. At 24 h post-infection, HO mice exhibited significantly higher bacterial loads than WT animals in both bladder and kidney (**Figure 2A&B**), indicating impaired early clearance of infection in mice with A-IC dysfunction. Assessment of multiple infection time points (24, 48, 72, and 96 h) showed that genotype-dependent differences were most pronounced at 24 h, whereas by 48 h post-infection, bladder CFUs were markedly reduced and no longer different in both genotypes, and kidney CFUs were undetectable (**Figure 2C–D**). Together, these findings indicate that A-IC dysfunction delays early bacterial clearance during UTI but does not impair eventual infection resolution in mice.

**Figure 2.**
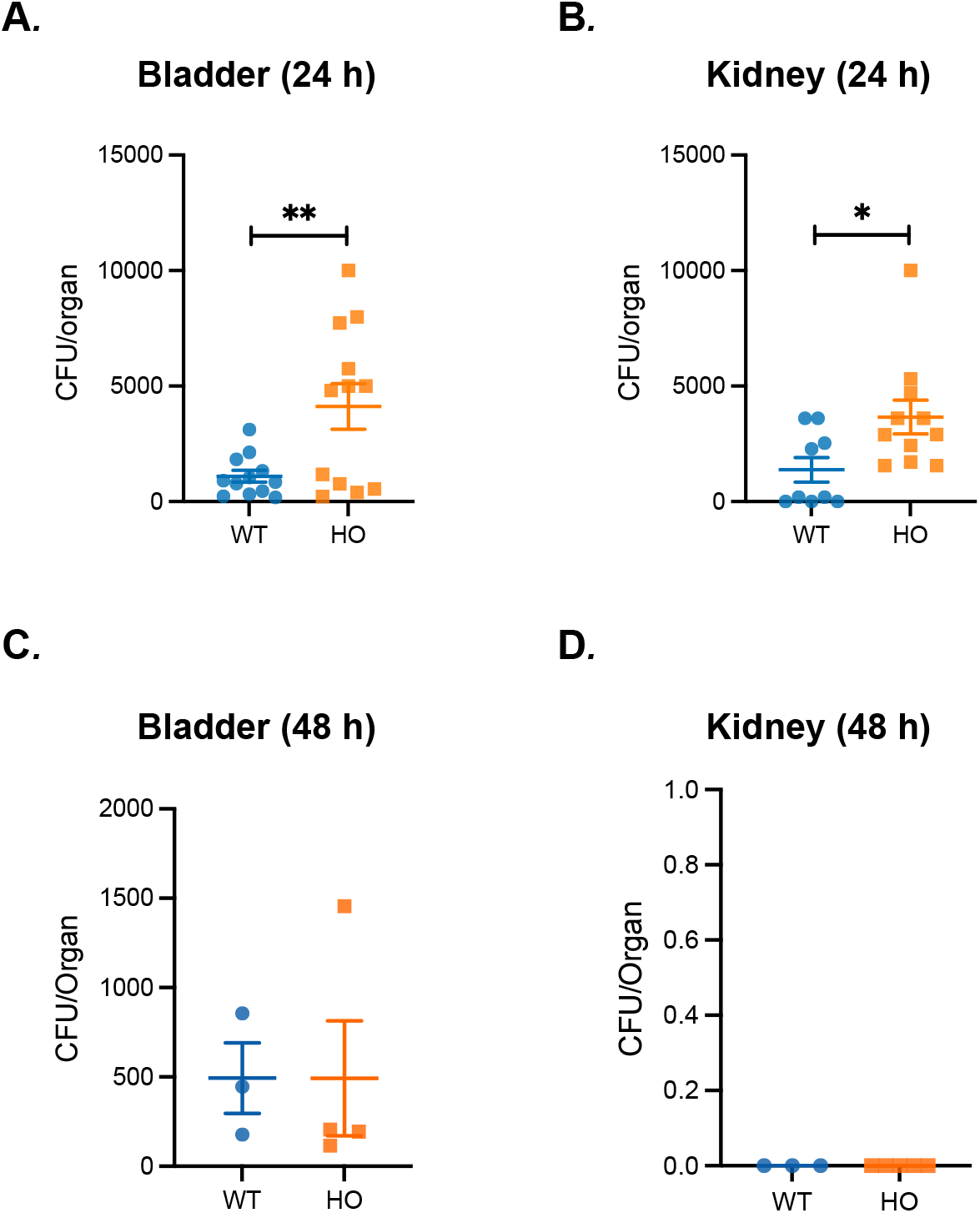
Bacterial burden in bladder and kidney at 24 h and 48 h after UPEC infection. CFUs were quantified in bladder (A&C) and kidney (B&D) homogenates from WT and HO mice at 24 h (A–B) and 48 h (C–D) post-infection. At 48 h, kidney CFUs were undetectable in all mice. Data are mean ± SEM; each point represents one animal. Statistical tests were unpaired *t*-test (24 h bladder) and Mann–Whitney test (24 h kidney; 48 h bladder). Kidney CFUs at 48 h were not analyzed due to all values being below detection limits (\**p* < 0.05; \*\**p* < 0.01).

### A-IC dysfunction alters AMP expression in dRTA kidneys

ICs contribute to renal antimicrobial defense through pathogen sensing, phagocytic activity, and AMP production ^12,14^. A-ICs, in particular, are as an important source of specific AMPs such as lipocalin-2 (LCN2) ^14^. We therefore examined whether A-IC dysfunction in the Ae1 R607H KI mice altered epithelial AMP expression during infection^12,14^.

Kidneys collected 24 h after UPEC or PBS inoculation were analyzed for AMP gene expression by qRT-PCR. *Lcn2*, galectin-3 (*Lgals3*), and cathelicidin-related antimicrobial peptide (*Camp*) transcripts were significantly increased in HO mice exposed to UPEC compared with their respective controls (**Figure 3**). In contrast, UPEC infection did not significantly alter *Lcn2, Lgals3*, or *Camp* mRNA expression in WT kidneys at 24 h. Therefore, although A-ICs are an important source of AMPs, their impairment in HO mice did not prevent induction of certain AMP transcripts during infection. In contrast, mRNA expression of ribonuclease A family member 4 (*Rnase4*), defensin beta 1 (*Defb1*), and adrenomedullin (*Adm*) showed no significant changes among groups (**Supplementary Figure 1A**). Protein levels of LCN2 measured by U-PLEX showed a modest, non-significant increase after UPEC infection in both WT and HO kidneys; however, only the HO group exhibited a trend consistent with its transcriptional upregulation (**Supplementary Figure 1B**). Together, these findings indicate that A-IC dysfunction is associated with altered expression of a subset of AMP transcripts during UTI in HO kidneys.

**Figure 3.**
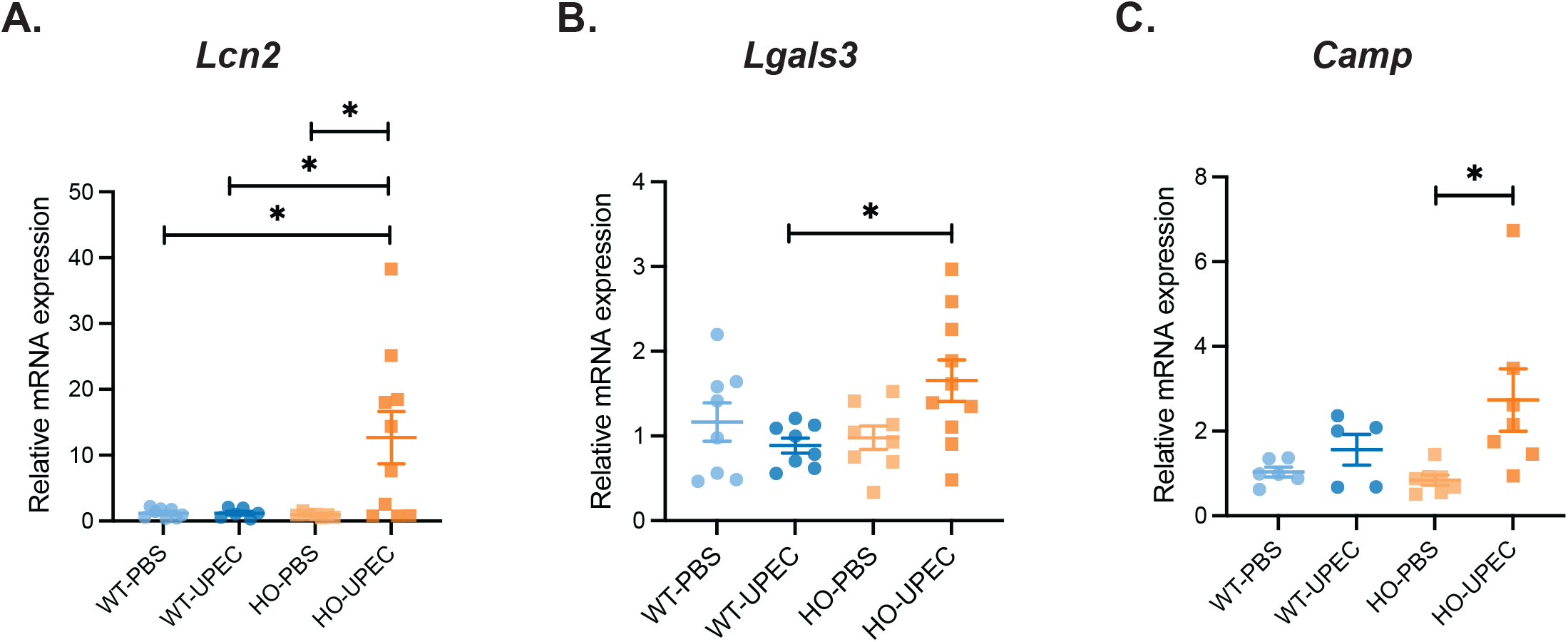
AMP gene expression in WT and HO kidneys following UPEC infection. qRT-PCR analysis of *Lcn2* (A), *Lgals3* (B), and *Camp* (C) mRNA expression in kidneys 24 h after PBS or UPEC inoculation. Data are presented as relative mRNA expression normalized to *Rplp0* and expressed relative to WT kidneys exposed to PBS using the 2^-ΔΔCT method. Data represent mean ± SEM, and each symbol indicates one animal. Statistical significance was determined using two-way ANOVA followed by Tukey’s multiple-comparisons test (\**p* < 0.05).

### A-IC dysfunction amplifies pro-inflammatory cytokine responses during UPEC infection

ICs not only serve as antimicrobial effectors but also contribute to local immune signaling through cytokine production and pH modulation in the CD ^12,15,18^. As A-IC dysfunction may alter the epithelial–immune communication, we next examined whether cytokine responses during infection differed between WT and HO mice. qRT-PCR analysis of kidneys collected 24 h post-infection revealed that interleukin-6 (*Il6*) and interleukin-1β (*Il1b*) transcripts were significantly elevated in HO UPEC mice compared with all other groups (**Figure 4A**). In contrast, UPEC infection did not significantly alter *Il6* or *Il1b* expression in WT kidneys at this time point.

**Figure 4.**
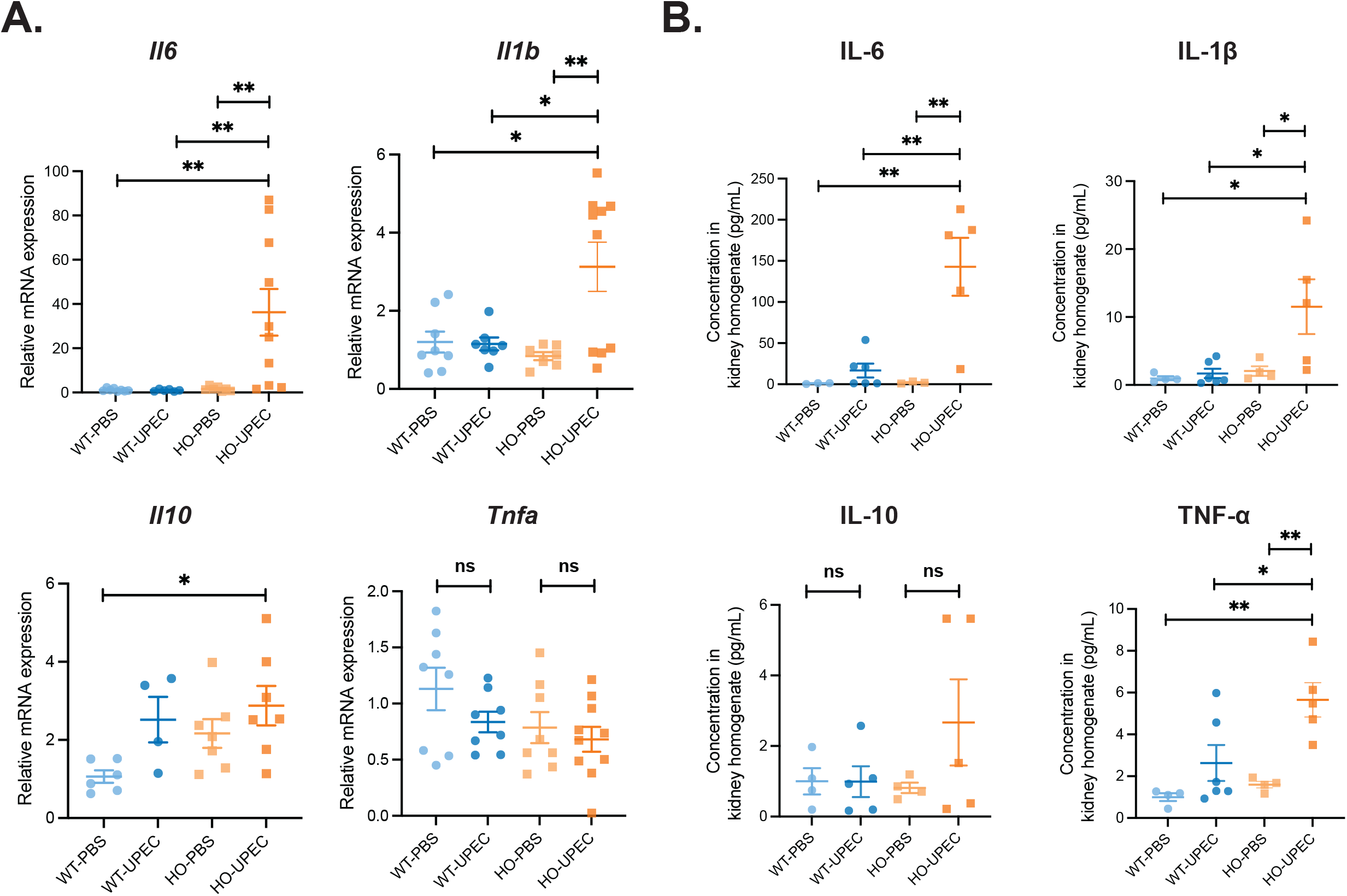
Cytokine expression in WT and HO kidneys following UPEC infection. (A) qRT-PCR analysis of *Il6, Il1b, Il10*, and *Tnfa* mRNA expression in kidneys from WT and HO mice 24 h after PBS or UPEC inoculation. Data are presented as relative mRNA expression normalized to *Rplp0* and expressed relative to WT PBS using the 2^-ΔΔCT method. (B) Protein abundance of IL-6, IL-1β, IL-10, and TNF-α measured by U-PLEX in kidneys collected 24 h after infection normalized to WT kidneys exposed to PBS. Data represent mean ± SEM, and each symbol indicates one animal. Statistical significance was determined by two-way ANOVA with Tukey’s multiple-comparisons test (\**p* < 0.05; \*\**p* < 0.01; ns, not significant).

Interleukin-10 (*Il10*) expression showed a modest but significant increase in HO UPEC kidneys only relative to WT PBS, while tumor necrosis factor-α *Tnfa* transcript levels showed no significant differences between PBS and UPEC within either genotype.

At the protein level, U-PLEX measurements generally mirrored the transcriptional data (**Figure 4B**). Interleukin-6 (IL-6) and interleukin-1β (IL-1β) concentrations were markedly higher in HO UPEC kidneys than in all other groups, and TNF-α levels were likewise increased in HO UPEC samples. Interleukin-10 (IL-10) protein abundance did not differ significantly between PBS and UPEC within either genotype, despite a trend for increase in HO UPEC samples. Together, these findings indicate that A-IC dysfunction is associated with an amplified pro-inflammatory cytokine response during UPEC infection.

### A-IC dysfunction enhances chemokine expression during UPEC infection

Cytokines and chemokines act together to coordinate immune cell recruitment during infection, and CD epithelial cells—including ICs—participate in renal immune signaling and chemokine responses under inflammatory conditions^15,34^. C-X-C motif chemokine ligand 2 (CXCL2) and C-C motif chemokine ligand 2 (CCL2) play essential roles in recruiting neutrophils and monocytes/macrophages to sites of renal injury or infection ^35,36^. Given the elevated cytokine response observed in HO UPEC mice, we next investigated whether chemokine expression was also altered during UPEC infection. qRT-PCR analysis revealed that *Cxcl2* and *Ccl2* transcript levels were significantly increased in HO UPEC kidneys compared with all other groups (**Figure 5**). In contrast, UPEC infection did not significantly alter the expression of these chemokines in WT kidneys. Together, these findings indicate that A-IC dysfunction is associated with increased chemokine expression during UPEC infection.

**Figure 5.**
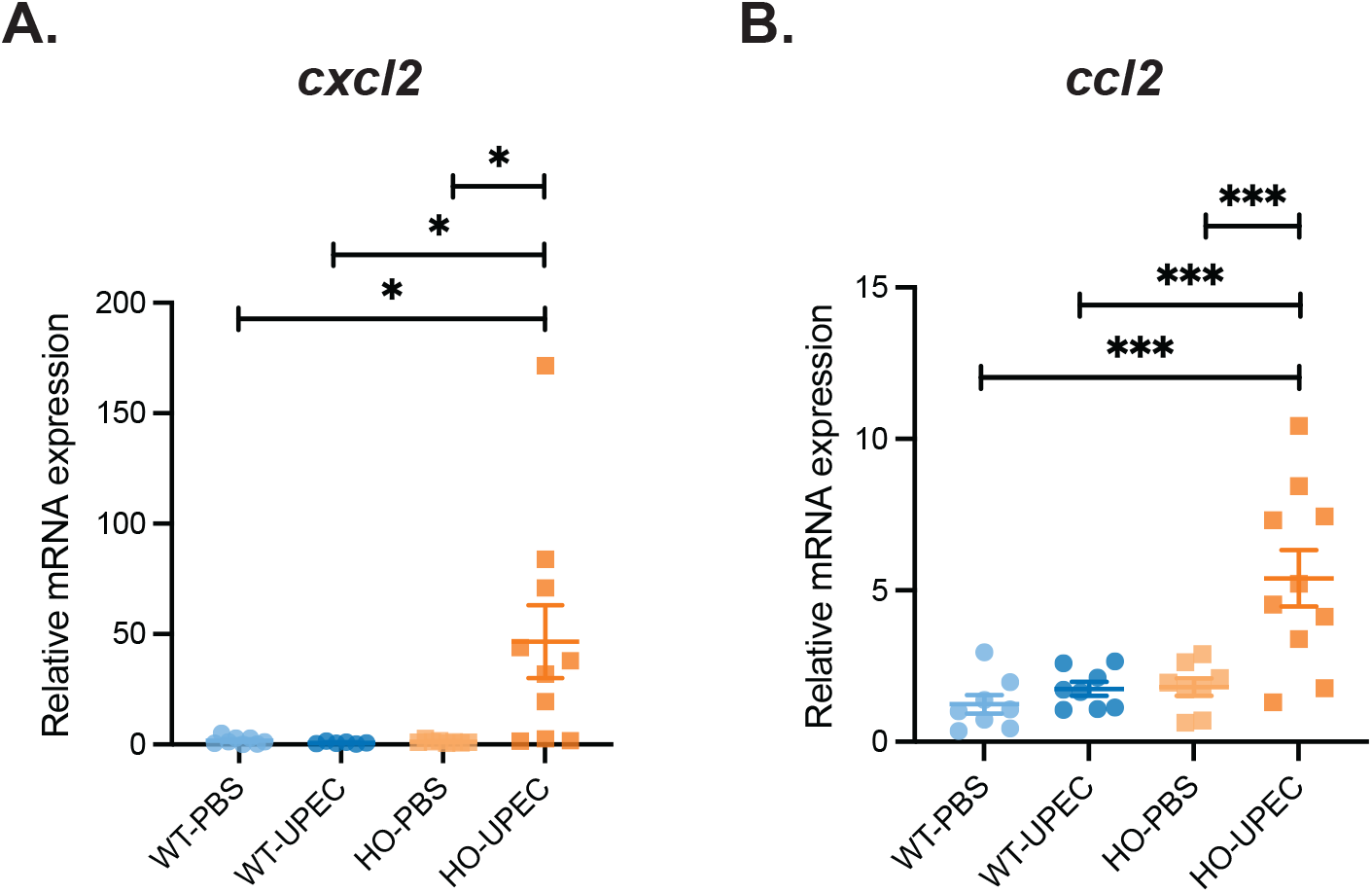
Chemokine expression in WT and HO kidneys following UPEC infection. mRNA levels of *Cxcl2* (A) and *Ccl2* (B) were measured by qRT-PCR and compared in kidneys from WT and HO mice 24 h after PBS or UPEC inoculation. Data are presented as relative mRNA expression normalized to *Rplp0* and expressed relative to WT PBS using the 2^-ΔΔCT method. Data represent mean ± SEM, and each symbol indicates one animal. Statistical significance was determined by two-way ANOVA with Tukey’s multiple-comparisons test (\**p* < 0.05; \*\*\**p* < 0.001).

### A-IC dysfunction disrupts immune cell recruitment during UPEC infection

Chemokines such as CXCL2 and CCL2 orchestrate the recruitment of neutrophils, monocytes, and macrophages during infection ^34^. Because ICs can modulate local immune cell trafficking through cytokine and chemokine signaling and given the marked increase in these chemokines in HO kidneys, we next examined whether A-IC dysfunction altered early leukocyte dynamics in the kidney.

To assess this, we quantified leukocyte populations 24 h post-infection by flow cytometry using the gating strategy shown in **Figure 6A**. The proportion of Cluster of Differentiation 45 (CD45^+^) leukocytes was significantly higher in HO kidneys exposed to UPEC compared with HO PBS, indicating enhanced inflammatory cell accumulation (**Figure 6B**). Notably, HO mice injected with PBS also displayed significantly fewer CD45^+^ cells than WT controls, suggesting baseline alterations in immune cell homeostasis. Analysis of specific myeloid subsets revealed that Lymphocyte antigen 6 complex locus G6D (Ly6G^+^) neutrophils were significantly elevated in HO kidneys following UPEC infection compared with WT (**Figure 6C**). WT UPEC kidneys also showed a significant increase in Lymphocyte antigen 6 complex locus C1 (Ly6C^+^) monocytes relative to WT PBS and these cells were also slightly more abundant in HO kidneys under PBS conditions compared with WT PBS controls. In HO mice, UPEC infection resulted in a non-significant upward trend toward higher Ly6C^+^ monocyte levels compared with all other condition (**Figure 6D**).

**Figure 6.**
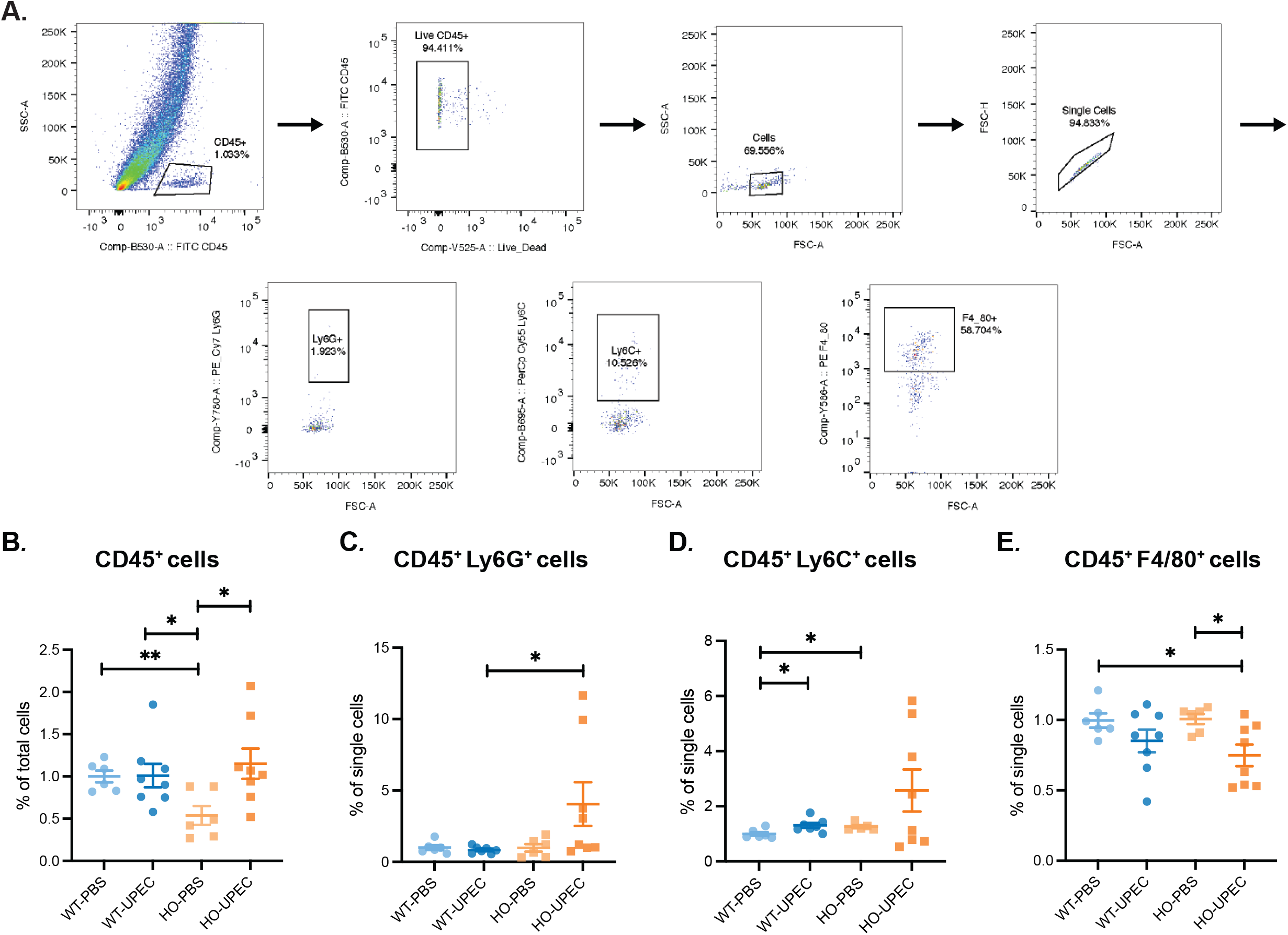
Gating strategy and quantification of renal leukocyte subsets 24h after UPEC infection. (A) Representative flow cytometry gating strategy for kidney leukocytes, showing sequential gating of CD45^+^ cells, exclusion of dead cells, debris removal (FSC-A vs SSC-A), singlet selection (FSC-A vs FSC-H), and downstream gating of CD45^+^ Ly6G^+^, CD45^+^ Ly6C^+^, and CD45^+^ F4/80^+^ cell subsets. (B–E) Quantification of CD45^+^ cells (B), CD45^+^ Ly6G^+^ cells (C), CD45^+^ Ly6C^+^ cells (D), and CD45^+^ F4/80^+^ cells (E) in kidneys 24 h after PBS or UPEC inoculation. Data are presented as mean ± SEM; each symbol represents one animal. Statistical significance was assessed using an unpaired two-tailed *t*-test for CD45^+^, Ly6C^+^, and F4/80^+^ subsets, and the Mann–Whitney test for the Ly6G^+^ subset (\**p* < 0.05; \*\**p* < 0.01).

In contrast, the proportion of EGF-like module-containing mucin-like hormone receptor-like 1 (F4/80^+^) cells was reduced in HO kidneys after UPEC infection compared with both HO PBS and WT PBS (**Figure 6E**). Because F4/80 is expressed by multiple myeloid populations—including resident macrophages and infiltrating monocytes—the observed reduction likely reflects an altered myeloid cell composition at this early time point (24 h) ^37^.

To further evaluate myeloid cell distribution, Cluster of Differentiation 68 (CD68) immunofluorescence was used as an additional marker of monocyte/macrophage populations ^38^ (**Figure 7A**). WT kidneys showed robust CD68^+^ cell presence under both PBS and UPEC conditions, whereas HO kidneys exposed to UPEC exhibited a visibly reduced CD68^+^ cell population. Quantification of CD68^+^/DAPI^+^ area ratios confirmed a significant decrease in CD68^+^ cell abundance in HO UPEC kidneys relative to other groups (**Figure 7B**). Because CD68 labels both macrophages and monocytes, these data collectively suggest that A-IC dysfunction alters early myeloid cell distribution rather than impairing a single cell population.

**Figure 7.**
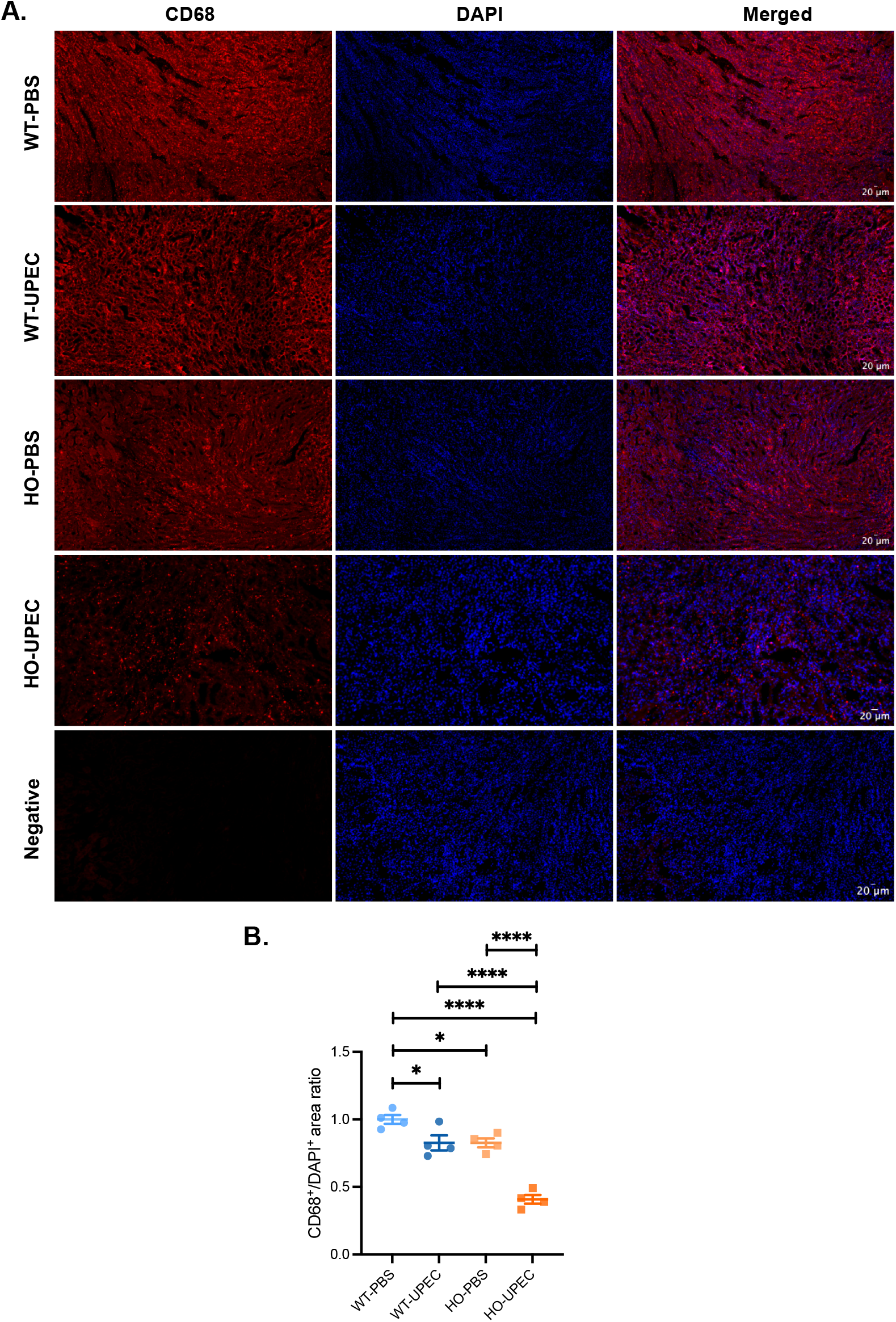
CD68^+^ myeloid cell distribution in WT and HO kidneys following UPEC infection. (A) Representative immunofluorescence images of kidney sections from WT and HO mice inoculated with PBS or UPEC. CD68^+^ myeloid-associated cells are shown in red, nuclei (DAPI) in blue, and merged images are displayed in the right column. The bottom row shows negative control images from kidneys that were not incubated with the primary antibody. All CD68 images are displayed using identical intensity settings. Scale bar = 20 µm. (B) Quantification of CD68^+^/DAPI^+^ area ratio in kidney sections, normalized to WT kidneys exposed to PBS. Each data point represents one animal. Statistical significance was determined by two-way ANOVA with Tukey’s multiple-comparisons test. Data are presented as mean ± SEM (\**p* < 0.05, \*\*\*\**p* < 0.0001).

Together, these findings indicate that A-IC dysfunction reshapes early leukocyte recruitment during UPEC infection—promoting exaggerated neutrophil accumulation while altering monocyte/macrophage-associated myeloid signatures—consistent with disrupted epithelial– immune communication in HO kidneys.

## DISCUSSION

This study reveals a crucial role for ICs in coordinating renal innate immune defense during UTI. Using the Ae1 R607H KI mouse model, which mirrors the dominant R589H dRTA *SLC4A1* variant, we show that A-IC dysfunction results in delayed bacterial clearance and exaggerated AMP, cytokine, and chemokine responses accompanied by altered renal myeloid-cell recruitment and inflammatory dynamics. These findings demonstrate that IC integrity is essential for maintaining balanced epithelial–immune communication during early infection in mice.

As previously reported by Mumtaz et al., A-IC depletion in Ae1 R607H KI mouse model predominantly occurs in the cortex, whereas medullary A-ICs remain present but exhibit reduced AE1 and V-ATPase B1 expression ^27^. In agreement with this characterization, our qRT–PCR analysis confirmed reduced expression of A-IC markers (*Slc4a1, Atp6v1b1*, and *Car2*) in HO kidneys under both PBS and UPEC conditions. UPEC infection did not alter expression of these markers, indicating that the underlying IC phenotype remains stable during early infection. Together, these observations confirm that this model faithfully recapitulates the molecular and cellular features of A-IC dysfunction and provide a unique opportunity for examining its immune phenotype.

ICs have traditionally been viewed as acid–base regulators, yet growing evidence indicates that they also contribute to renal innate immunity through expression of pattern-recognition receptors and secretion of antimicrobial and inflammatory mediators ^12,15,39^. Consistent with this broader role, we observed significantly higher bacterial loads in the bladder and kidney of HO mice 24 h after UPEC infection. These differences resolved by 48 h, consistent with the normal trajectory of bacterial clearance in standard murine UTI models ^30,40^. The increased early bacterial burden in HO mice likely reflects the combined effects of reduced A-IC function and impaired urinary acidification, both of which can diminish early epithelial antimicrobial defenses and facilitate initial UPEC expansion ^27,41,42^.

At the molecular level, A-IC dysfunction was associated with significant changes in the renal antimicrobial landscape. *Lcn2, Lgals3*, and *Camp* transcripts were markedly elevated in HO kidneys following UPEC infection, whereas *Rnase4, Defb1*, and *Adm* remained unchanged. The upregulation of these AMPs despite reduced A-IC function suggests compensatory activation by other epithelial or immune cell populations, consistent with prior reports of Lcn2 induction in distal nephron epithelial cells during infection or ischemic injury ^14,43^. Notably, WT kidneys did not show robust AMP induction at 24 h, likely because intact ICs and effective urine acidification restrict early bacterial expansion and keep epithelial stress below the threshold for AMP transcriptional activation. In HO mice, impaired urinary acidification and higher early bacterial burden generate a stronger inflammatory and epithelial stress, driving AMP transcription yet not ensuring effective antimicrobial function. However, because many AMPs exhibit pH-dependent activity, their effectiveness may be limited in the alkaline luminal environment characteristic of dRTA, meaning that increased AMP expression may not translate into efficient bacterial clearance in vivo ^27,41,42^.

Analysis of cytokine profiles revealed that A-IC dysfunction is associated with an amplified pro-inflammatory response during UPEC infection, with elevated *Il6* and *Il1b* transcripts and corresponding proteins, and significantly elevated TNF-α protein in HO UPEC kidneys, whereas WT kidneys did not show induction of these cytokines at 24 h. *Il10* transcripts also showed a modest but significant increase in HO UPEC kidneys, although this induction did not translate into higher IL-10 protein, suggesting post-transcriptional regulation or delayed translation. Together, these findings indicate that HO kidneys develop a cytokine milieu dominated by pro-inflammatory IL-6, IL-1β, and TNF-α, with insufficient early anti-inflammatory compensation despite *Il10* transcriptional induction. Robust IL-6 and IL-1β induction has also been documented in UPEC infection models using C3H (HeN and HeOuJ) and C57BL/6 mice, where early cytokine and chemokine induction correlates with neutrophil recruitment and contributes to bacterial clearance ^44–46^. In our study, a comparable cytokine increase occurred only in HO mice, indicating that A-IC dysfunction lowers the threshold for epithelial and immune activation during infection. This finding indicates that under dRTA conditions, impaired IC function does not only diminish epithelial defenses but also predisposes the kidney to amplified cytokine signaling. Because ICs contribute to renal innate immune signaling, their reduced abundance or altered function in HO mice likely removes a stabilizing epithelial checkpoint that normally helps limit the magnitude of early cytokine responses ^7^. The result is a more pronounced inflammatory milieu in HO kidneys during the initial phase of infection.

In line with the lack of AMP induction in infected WT UPEC mice, the lack of cytokine upregulation in WT kidneys at 24 h likely reflects more effective early bacterial control in WT mice. Intact IC function and normal urinary acidification likely reduce epithelial stress and prevent the strong inflammatory signaling observed in HO mice. Moreover, innate immune responses during ascending UTI can occur rapidly in WT mice, with epithelial and leukocyte activation detected within hours of infection in murine models ^47^. Thus, WT mice may have generated a transient cytokine surge that has already begun to resolve by 24 h post-infection. This interpretation is further supported by Hamilton and colleagues’ work who observed minimal IL-6, IL-1β, and TNF-α at 24 h despite confirmed infection, indicating that cytokine elevations at this time point may be transient or low-level ^48^.

Consistent with cooperative function of cytokines and chemokines ^34^, the strong cytokine response in HO mice was accompanied by marked upregulation of *Cxcl2* and *Ccl2*, key mediators of neutrophil and monocyte recruitment. Comparable induction of these chemokines has been reported in UPEC-infected C3H and C57BL/6 mouse models, where *Cxcl2* and *Ccl2* drive early leukocyte infiltration and contribute to bacterial clearance ^46,49^. ICs also upregulate chemokines such as Cxcl2 and Ccl2 during epithelial stress ^19^. Our findings show that in the context of dRTA, impaired A-IC lead to amplified chemokine responses, suggesting a role in fine-tuning epithelial– immune communication within the CD microenvironment.

Flow-cytometric profiling revealed that the amplified cytokine and chemokine responses in HO mice were accompanied by marked alterations in early intra-renal myeloid-cell composition. Total CD45^+^ leukocytes, Ly6G^+^ neutrophils and Ly6C^+^ monocytes were significantly increased in HO kidneys following UPEC infection, indicating exaggerated neutrophil and monocyte recruitment. In contrast, the proportion of F4/80^+^ cells was reduced in HO kidneys at 24 h. Because F4/80 is expressed by multiple myeloid populations—including resident macrophages and monocyte-derived cells—this reduction could reflect an altered myeloid composition rather than a selective loss of macrophages at this early time point ^37^. Immunofluorescence analysis using CD68, another marker of monocyte/macrophage populations, further supported this conclusion, with robust CD68^+^ signal in WT kidneys under both PBS and UPEC conditions, but a significant reduction in CD68^+^ staining in infected HO kidneys ^38^. Taken together, these findings indicate that A-IC dysfunction does not prevent monocyte entry into the kidney but disrupts the balance of myeloid subsets, favoring exaggerated neutrophil accumulation while limiting the early expansion or retention of monocyte-derived macrophage populations.

The Ae1 R607H KI model recapitulates key features of dominant *SLC4A1*-mediated dRTA while preserving overall kidney architecture, enabling focused investigation of IC-dependent aspects of epithelial–immune interactions. A limitation of our study is the focus on a single early time point (24 h), which restricts interpretation to initial stages of infection. Future studies incorporating longitudinal sampling, flow cytometric profiling across later phases, and IC-specific transcriptomic analyses will be important for defining how regional IC dysfunction shapes cytokine, chemokine, and myeloid-cell responses over the course of infection.

In conclusion, our study demonstrates that A-IC dysfunction is an important modifier of early renal immune responses during UTI. A loss of A-IC phenotype—together with impaired urinary acidification—was associated with altered antimicrobial, cytokine, and chemokine responses and with an early myeloid-cell imbalance characterized by heightened neutrophil recruitment. These findings suggest that IC function contributes to the regulation of epithelial–immune communication in the CD and highlight the importance of intact IC physiology regulation for maintaining effective antimicrobial defense in the distal nephron.

## ACKNOWLEDGEMENTS

The authors want to thank Drs. Sylvie Breton (Université Laval), Jean-Francois Cailhier and Patrick Laplante (Centre Hospitalier de l’Université de Montréal) for their help in the immunostaining, flow cytometry experiments and for helpful discussions; the authors also gratefully acknowledge Drs. Aja Rieger, Lai Xu, and Rikus Niemand from the Flow Cytometry Facility, as well as Dr. Hilmar Strickfaden and Kiera Smith from the Cell Imaging Facility at the University of Alberta, for their valuable technical assistance. The University of Alberta Advanced Cell Exploration Core Facility (RRID:SCR_019182) receives financial support from the Faculty of Medicine & Dentistry, the University Hospital Foundation, Striving for Pandemic Preparedness – The Alberta Research Consortium, and Canada Foundation for Innovation awards. The UPEC strain used in this study was kindly provided by Dr. Gregory Tyrrell (Alberta Precision Laboratories). FC was supported by Graduate Recruitment Scholarship and a Graduate Student Award from the University of Alberta. GE received Graduate Student Engagement Scholarships, Faculty of Medicine and Dentistry Delnor Scholarship, Faculty of Medicine and Dentistry 75^th^ Anniversary award and an Alberta Graduate Excellence Scholarship. This study was supported by funding from the Natural Sciences and Engineering Research Council (Discovery Grant RGPIN-2017-06432), Canadian Institutes of Health Research (PJT-168871) and the Kidney Foundation of Canada (2020 KHRG-666615 & 25KHRG-1422220).

---

**Figure S1.**
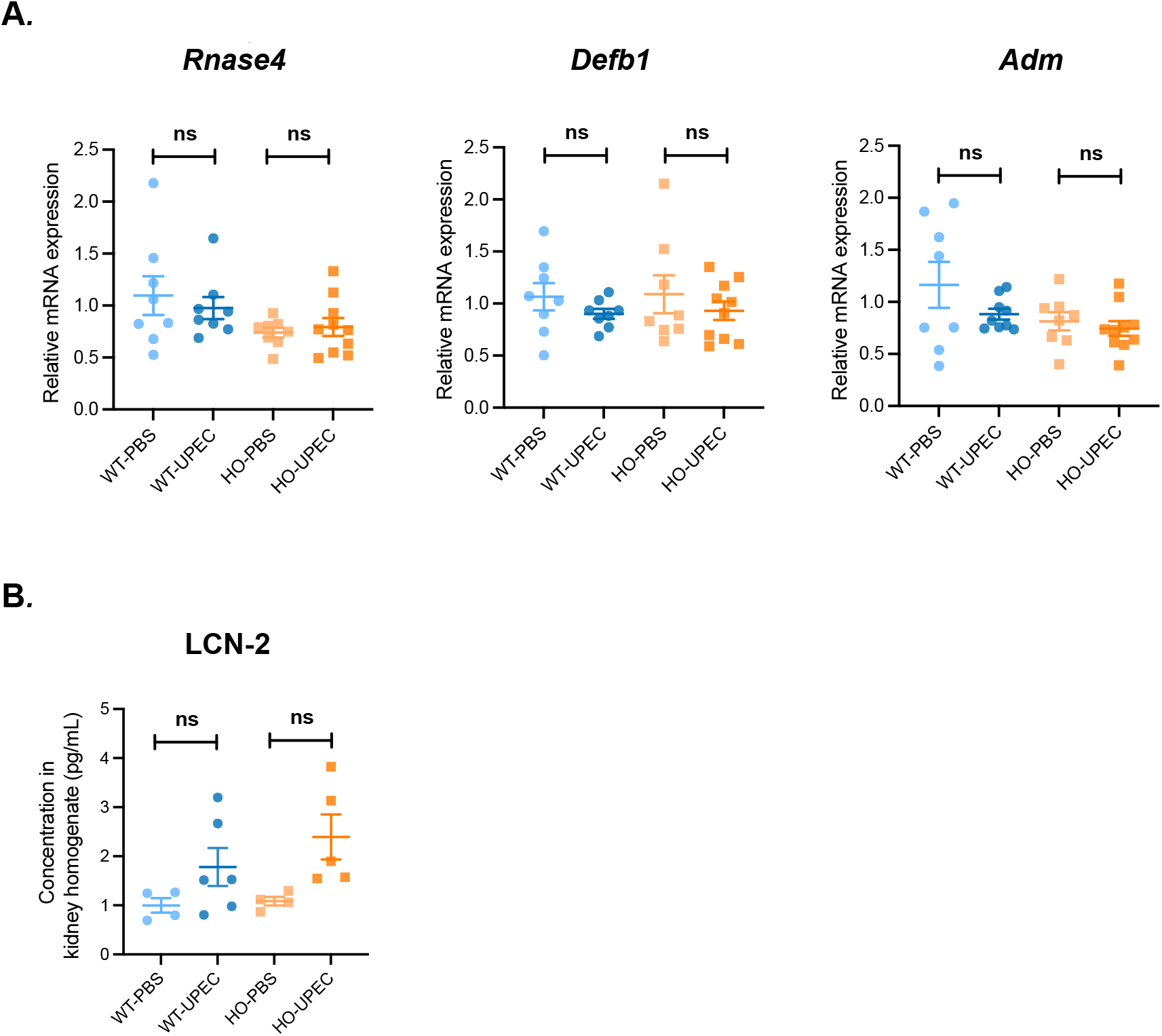

