## Supplementary Material for "A-Intercalated Cell Dysfunction Disrupts Renal Epithelial–Immune Balance and Impairs Host Defense During UTI"

### Supplementary Methods

#### 1. Animal Model

All animal procedures were conducted in accordance with the guidelines of the Canadian Council on Animal Care (CCAC) and were approved by the Animal Care and Use Committee (ACUC) at the University of Alberta under Animal Use Protocol (AUP No. 1277). Female wild-type (WT) and Ae1 R607H homozygous (HO) dRTA KI mice aged 8–10 weeks were used for all experiments. The Ae1 R607H mice have previously been characterized [1].All animals were housed in Health Sciences Laboratory Animal Services (HSLAS) Biosafety Level 2 under a 12-hour light/dark cycle with *ad libitum* access to water and standard chow (PicoLab Mouse Diet 20, Cat. no. 5058).

#### 2. Bacterial Preparation

A Gram-negative, non-haemolytic uropathogenic *Escherichia coli* (UPEC) strain, originally isolated from a female patient with pyelonephritis, was kindly provided by Dr. Gregory Tyrell (Alberta Precision Laboratory) and used for this study. Fresh bacteria were prepared according to a previously published protocol [2]. Bacteria from the stock tube were streaked onto Luria-Bertani (LB) agar plates and incubated overnight at 37 °C. A single colony was then inoculated into 5 mL of LB broth and grown in a shaking incubator at 200 rpm and 37 °C for 24 hours. Subsequently, 500 µL of this overnight culture was sub-cultured into 5 mL of fresh LB broth and incubated under the same conditions for an additional 24 hours. Bacterial cells were then harvested by centrifugation at 3000 × g for 10 minutes, and the resulting pellet was resuspended in 1–5 mL of sterile phosphate-buffered saline (PBS), depending on the experiment. A 1:5000 dilution was plated on LB agar and incubated overnight at 37 °C to determine colony-forming units (CFUs). This allowed calculation of the bacterial concentration, typically yielding 20–100 × $10^6$ CFUs per 50 µL, to ensure consistency across experiments.

#### 3. UTI Induction

Mice were deprived of drinking water for 30 minutes prior to anesthesia to minimize immediate voiding post-inoculation. Before anesthesia, mice were placed in a sterile container, and gentle massage of the caudal abdomen was performed to induce urination and ensure an empty bladder [3]. Mice were then anesthetized with isoflurane, and upon reaching a surgical plane of anesthesia, the vulva and perineum were swabbed with 70% ethanol to reduce the risk of external contamination. A sterile 1 cc syringe attached to a lubricated, sterile catheter/cannula was inserted into the urethra, and 50 µL of either bacterial suspension (20–100 × $10^6$ colony-forming units (CFUs)) or sterile phosphate-buffered saline (PBS) was slowly instilled into the bladder [2]. Mice were maintained under anesthesia for up to 20 minutes post-inoculation to prevent immediate voiding, then allowed to recover on a warming pad under close observation before being returned to their home cages.

#### 4. Tissue Collection and Bacterial Load Quantification

Mice were euthanized 24 h post-infection via intraperitoneal injection of pentobarbital (50 mg/kg), followed by perfusion with PBS containing 1 unit/mL heparin to remove circulating blood prior to tissue collection. Bladders and kidneys were collected aseptically. Half of the bladder and half of one kidney were homogenized in 200 µL of sterile PBS using a tissue homogenizer. The entire homogenate was plated onto LB agar and incubated overnight at 37 °C for CFU enumeration.

#### 5. RNA Isolation and Quantitative Reverse Transcription Polymerase Chain Reaction (qRT-PCR)

Half of a kidney was placed in RNAlater (Thermo Fisher Scientific Cat. no. 00712711) immediately after collection and later stored at −80 °C. Total RNA was extracted using TRIzol reagent (Invitrogen Cat. no. 15596018) following the manufacturer's instructions. RNA yield was determined using a NanoDrop™ 2000C spectrophotometer (Thermo Fisher Scientific). Reverse transcription was performed using 2 μg of RNA with either SuperScript™ II (Invitrogen Cat. no. 100004925) or the High Capacity cDNA Reverse Transcription Kit (Thermo Fisher Scientific Cat. no. 4368814), following the respective manufacturer's protocols. qRT-PCR was conducted in triplicate for each cDNA sample using TaqMan PCR Master Mix on a QuantStudio 6 Pro qRT-PCR system (Thermo Fisher Scientific). Gene expression was analyzed using the $2^{-\Delta\Delta Ct}$ method [4]. mRNA levels of target genes were normalized to the housekeeping gene ribosomal protein lateral stalk subunit P0 (*Rplp0*). The lack of variation in *Rplp0* gene expression between experimental conditions was verified. The primer sequences used for each gene are provided in **Table S1**.

#### 6. Protein Extraction and Quantification by U-PLEX Assay

One half of a freshly dissected kidney was decapsulated and mechanically homogenized in ice-cold RIPA lysis buffer (5 mM EDTA pH 8.0, 150 mM NaCl, 0.5% Na-Deoxycholate, 1% Nonidet P-40 (NP-40), 0.1% SDS, 50 mM Tris), supplemented with protease inhibitors (Complete Mini, EDTA-free, Roche) and phosphatase inhibitors (PhosSTOP, Roche). The homogenate was incubated on ice for 1 hour with vortexing every 15 minutes, followed by centrifugation at 18,407 × g for 15 minutes at 4 °C. The resulting supernatant was collected and stored at –80 °C until protein quantification.

Protein levels in the lysates were quantified using a custom-designed U-PLEX multiplex immunoassay (Meso Scale Discovery, MSD, Cat. no. K15069L-1) targeting selected cytokines and antimicrobial proteins, according to the manufacturer's instructions. Standards and appropriately diluted samples were incubated with pre-coated 96-well plates for 1 hour at room temperature with shaking. After washing, SULFO-TAG-conjugated detection antibodies were added and incubated for an additional hour. Plates were read using the MESO QuickPlex SQ 120 reader (MSD) at the University of Alberta Advanced Cell Exploration Core facility (RRID:SCR_019182), and data were analyzed using Discovery Workbench software (v4.0). All samples were run in duplicate, and concentrations were calculated based on standard curves.

7. **Flow Cytometry**

Following euthanasia and perfusion as described above, both kidneys were collected and cut into 3–4 pieces. Single-cell suspensions were prepared using the Multi-Tissue Dissociation Kit 1 (Miltenyi Biotec, Cat. no. 130-110-201) and gentleMACS™ Octo Dissociator with Heaters (Cat. no. 130-096-427) according to the manufacturer's instructions. Erythrocytes were lysed briefly with Ammonium Chloride Potassium (ACK) lysis buffer (0.15 M $NH_4Cl$, 1.0 mM $KHCO_3$, and 100 mM EDTA) and resuspended in fluorescence-activated cell sorting (FACS) buffer (PBS with 1% FBS and 2 mM EDTA). After final filtration through 40 μm strainers (Fisherbrand™, Cat. no. 22363547), cells were counted using a hemocytometer and adjusted to the appropriate concentration for antibody staining ($10^6$ cells per 100 μL). Single cell suspensions were incubated with Fc receptor blocking antibody at 4°C for 5 minutes to prevent nonspecific binding [4]. Cells were then stained with fluorochrome-conjugated antibodies and a live/dead viability dye (Aqua fluorescent reactive dye, Live/Dead™ Fixable Dead Cell Stain Kits, Thermofisher Scientific, Cat. no. L34957) according to manufacturer instructions. Compensation controls were prepared using UltraComp eBeads™ Plus Compensation Beads (Thermo Fisher Scientific Cat. no. 01-3333-41) stained individually with each antibody [5]. Fluorescence Minus One (FMO) controls were included for each fluorochrome to set gating boundaries. Instrument settings were optimized by running unstained (negative control) and fully stained samples to adjust photomultiplier tube (PMT) voltages. Data were acquired on a BD LSRFortessa X-20 using FACSDiva at the University of Alberta Faculty of Medicine & Dentistry Flow Cytometry Facility (RRID:SCR_019195); 30,000 events were collected per sample. Data were analyzed in FlowJo v10 under the University of Alberta institutional license. Compensation matrices were applied using single-stained controls, and gating strategies were guided by FMO controls to ensure accurate identification of cell populations. The gating strategy began with CD45 vs side scatter (SSC) to identify leukocytes, followed by CD45 vs Live/Dead staining to exclude non-viable cells. A FSC vs SSC gate was then used to remove debris, and single cells were selected using FSC-A vs FSC-H. Final gates were applied to define specific immune cell populations based on surface marker expression. Details of the antibodies used are provided in **Table S2**.

### 8. Immunofluorescence

After perfusion, kidneys were fixed in 4% paraformaldehyde (PFA) overnight at 4°C. The following day, tissues were cryoprotected by sequential incubation in 17.5 % sucrose for 2 hours, followed by 35 % sucrose overnight at 4°C. Kidneys were then embedded in Tissue-Tek® O.C.T. compound (Sakura, Cat. no. 4583), snap-frozen in liquid nitrogen, and stored at –80°C until further processing. Cryosections (10 µm thick) were mounted onto charged glass slides (ThermoFisher, Cat. no. 22-265446) and stored at –80°C for future analysis.

For immunostaining, cryosections were air-dried at room temperature (RT) for 3-5 minutes, fixed in 4% PFA for 20 min on a gentle shaker, and rinsed in PBS for 30 min. Sections were permeabilized with 0.1% Triton X-100 in PBS for 5 min and blocked with 4% bovine serum albumin (BSA) in PBS for 1 h at RT. Slides were incubated overnight at 4 °C with rat anti-mouse CD68 primary antibody (Bio-Rad, Cat. no. MCA1957GA) diluted in 2% BSA, washed 3×5 min in PBST (0.1 % Tween-20 in PBS), and incubated with goat anti-rat cyanine 3 secondary antibody (ThermoFisher, Cat. no. A10522) for 1 hour at RT. After two subsequent PBST washes, a third wash was performed using DAPI (Sigma-Aldrich, Cat. no. D9564) to stain nuclei. Slides were then mounted using DAKO Mounting Medium (DakoCytomation, Cat. no. S3022). Negative control sections processed without primary antibody were included to confirm staining specificity. Imaging was performed on a ZEISS Axio Scan.Z1 slide scanner using a 10× objective at the University of Alberta Faculty of Medicine & Dentistry Cell Imaging Core (RRID:SCR_019200). For analysis, $CD68^+$ and $DAPI^+$ areas were measured in FIJI (ImageJ2, version 2.14.0/1.54f) from four planes per section, averaged, and their ratio was calculated.

**9. Statistical Analysis**

Statistical analyses were performed using GraphPad Prism (v10.4.1). When appropriate, data were normalized to the average value of the WT PBS group before statistical testing. Data were assessed for normality, and outliers identified by the software were excluded. Depending on the distribution and experimental design, comparisons between two groups were made using either an unpaired two-tailed Student's *t*-test or the Mann–Whitney *U* test. For multiple group comparisons, two-way ANOVA followed by Tukey's multiple-comparisons test was used. For flow cytometry experiments where specific groups were directly compared, unpaired two-tailed *t*-tests were applied. A *p*-value of less than 0.05 was considered statistically significant. Results are expressed as mean ± standard error of the mean (S.E.M.).

All reagents, catalog numbers, and software versions are listed for reproducibility. Supplementary Tables S1 and S2 provide primer sequences and antibody panels, respectively.

**Supplementary Figures Legends**

**Supplementary Figure 1. Additional antimicrobial peptide expression in WT and HO kidneys.** (A) qRT-PCR analysis of *Rnase4*, *Defb1*, and *Adm* transcripts 24 h after PBS or UPEC inoculation. Data are presented as relative mRNA expression, normalized to *Rplp0* and expressed relative to WT kidneys exposed to PBS using the $2^{-\Delta\Delta CT}$ method. No significant differences were detected among groups. (B) LCN2 protein levels measured by U-PLEX in kidneys collected 24 h after infection, normalized to WT kidneys exposed to PBS. Data are mean ± SEM, and each symbol indicates one animal. Statistical significance was assessed using two-way ANOVA followed by Tukey's multiple-comparisons test **(**ns, not significant).

**Supplementary Tables**

**Table S1. Primer and probe sequences of mouse genes used for qRT-PCR analysis**

| **Gene** | **Assay Name** | **Forward Primer** | **Reverse Primer** | **Probe** |
|---|---|---|---|---|
| *Atp6v1b1* | N134157.1.pt.Atp6v1b1 | 5′-CTTCTCACACTGGTAGGCAAG-3′ | 5′-AGTCAGATTTTCGAGCAGAATGG-3′ | 5′-/56-FAM/CCGCTCAAT/ZEN/CGTAGGGTCATTGGC/3IABkFQ/-3′ |
| *Car2* | N009801.1.pt.Car2 | 5′-GGAGCAAGGGTCGAAGTTAG-3′ | 5′-GATGGATTGGCTGTTTTGGG-3′ | 5′-/56-FAM/CCGCACGCT/ZEN/TCCCCTTTGTTTT/3IABkFQ/-3′ |
| *Ccl2* | N011333.1.pt.Ccl2 | 5′-GCTCTCCAGCCTACTCATTG-3′ | 5′-GTCCCTGTCATGCTTCTGG-3′ | 5′-/56-FAM/TGCAGTTAA/ZEN/CGCCCCACTCACC/3IABkFQ/-3′ |
| *Cxcl2* | N009140.1.pt.Cxcl2 | 5′-CTTCCGTTGAGGGGACAGC-3′ | 5′-GAAGTCATAGCCACTCTCAAGG-3′ | 5′-/56-FAM/TCCTTTCCA/ZEN/GGTCAGTTAGCCTTGC/3IABkFQ/-3′ |

| | | | | |
|---|---|---|---|---|
| *Il1b* | N008361.1.pt.Il1b | 5′-TTCTCCACAGGCCACAAATGAG-3′ | 5′-ACGGACCCCAAAAGATGAAG-3′ | 5′-/56-FAM/AGAGCATCC/ZEN/AGCTTCAAATCTCGCA/3IABkFQ/-3′ |
| *Il6* | Mm.PT.58.13354106 | 5′-GATACCACTCCCAACAGACC-3′ | 5′-CAAGTGCATCATCGTTGTTCA-3′ | 5′-/56-FAM/CCATTGCAC/ZEN/AACTCTTTTCTCATTTCCACG/3IABkFQ/-3′ |
| *Lcn2* | Mm.PT.58.10167155 | 5′-CTACAATGTCACCTCCATCCTG-3′ | 5′-CCTGTGCATATTTCCCAGAGT-3′ | 5′-/56-FAM/TGTTCTGAT/ZEN/CCAGTAGCGACAGCC/3IABkFQ/-3′ |
| *Lgals3* | N010705.2.pt.Lgals3 | 5′-AGTTGGCTGATTTCCCGC-3′ | 5′-GCCTTCCCCTTTGAGAGTG-3′ | 5′-/56-FAM/AGTAGGTGA/ZEN/GCATCGTTGACCGC/3IABkFQ/-3′ |
| *Rplp0* | Mm.PT.58.43894205 | 5′-TTATAACCCTGAAGTGCTCGAC-3′ | 5′-CGCTTGTACCCATTGATGATG-3′ | 5′-/56-FAM/AGGCCCTGC/ZEN/ACTCTCGCTT/3IABkFQ/-3′ |
| *Slc4a1* | SLC4A1 (kAE1) | 5′-AAGAGGTCAAGGAACAGCG-3′ | 5′-GTACAGGAAGATGCCGAAGAG-3′ | 5′-/56-FAM/CCCACAAGC/ZEN/ACAGAGACCAGGAG/3IABkFQ/-3′ |

**Table S2. Primary Antibodies Used for Flow Cytometry Analysis**

| Name | Clone | Fluorochrome | Isotype | Supplier | Cat. no. |
|---|---|---|---|---|---|
| CD16/CD32 (Fc Block) | 2.4G2 | - | Rat (SD) IgG2b, κ | BD Pharmingen™ | 553141 |
| CD45 | 30-F11 | FITC | Rat (LOU) IgG2b, κ | BD Pharmingen™ | 553080 |
| Ly6G | 1A8 | PE/Cyanine7 | Rat IgG2a, κ | BioLegend® | 127617 |
| Ly6C | AL-21 | PerCP-Cy™5.5 | Rat IgG2a, κ | BD Pharmingen™ | 560525 |
| F4/80 | BM8 | PE | Rat IgG2a, κ | BioLegend® | 123109 |
